# Whole Genome Sequencing in Psychiatric Disorders: the WGSPD Consortium

**DOI:** 10.1101/160499

**Authors:** Stephan J. Sanders, Benjamin M. Neale, Hailiang Huang, Donna M. Werling, Joon-Yong An, Shan Dong, Whole Genome Sequencing for Psychiatric Disorders, Goncalo Abecasis, P. Alexander Arguello, John Blangero, Michael Boehnke, Mark J. Daly, Kevin Eggan, Daniel H. Geschwind, David C. Glahn, David B. Goldstein, Raquel E. Gur, Robert E. Handsaker, Steven A. McCarroll, Roel A. Ophoff, Aarno Palotie, Carlos N. Pato, Chiara Sabatti, Matthew W. State, A. Jeremy Willsey, Steven E. Hyman, Anjene M. Addington, Thomas Lehner, Nelson B. Freimer

**Author notes:** Please address correspondence to (S. E. H.), (A. M. A.), (T. L.), and (N. B. F.). These authors contributed equally to this work (Stephan J. Sanders, Benjamin M. Neale).

## Abstract

As technology advances, whole genome sequencing (WGS) is likely to supersede other genotyping technologies. The rate of this change depends on its relative cost and utility. Variants identified uniquely through WGS may reveal novel biological pathways underlying complex disorders and provide high-resolution insight into when, where, and in which cell type these pathways are affected. Alternatively, cheaper and less computationally intensive approaches may yield equivalent insights. Understanding the role of rare variants in the noncoding gene-regulating genome, through pilot WGS projects, will be critical to determine which of these two extremes best represents reality. With large cohorts, well-defined risk loci, and a compelling need to understand the underlying biology, psychiatric disorders have a role to play in this preliminary WGS assessment. The WGSPD consortium will integrate data for 18,000 individuals with psychiatric disorders, beginning with autism spectrum disorder, schizophrenia, bipolar disorder, and major depressive disorder, along with over 150,000 controls.

## Main text

Genetic variation is a major contributor to neuropsychiatric disorders. The variants responsible likely include the complete range of sizes, from single nucleotides to large structural variants, and the full spectrum of population frequency, from common variants to rare variants unique to a family or individual. For severe, early onset neuropsychiatric disorders, such as autism spectrum disorder (ASD) and schizophrenia, natural selection limits the population frequency of variants so that variants with larger effect sizes are extremely rare^1,2^. Over the past decade, genomic technologies have advanced our understanding of neuropsychiatric disorders, yet remaining limitations in technology and cohort sizes have limited progress in identifying inherited rare variants.

Genome-wide association studies (GWAS) using genotyping arrays have detected over 100 regions (loci) at which common genetic variants (population frequency ≥2%), are associated with a psychiatric diagnosis (Table 1). Individually, these variants exert small effects and thus require very large sample sizes for detection (Table 1). Common risk variants can provide a window into the molecular architecture of these disorders. For example, common variants suggest a previously unrecognized role for the complement cascade in schizophrenia^3^.

**Table 1.**
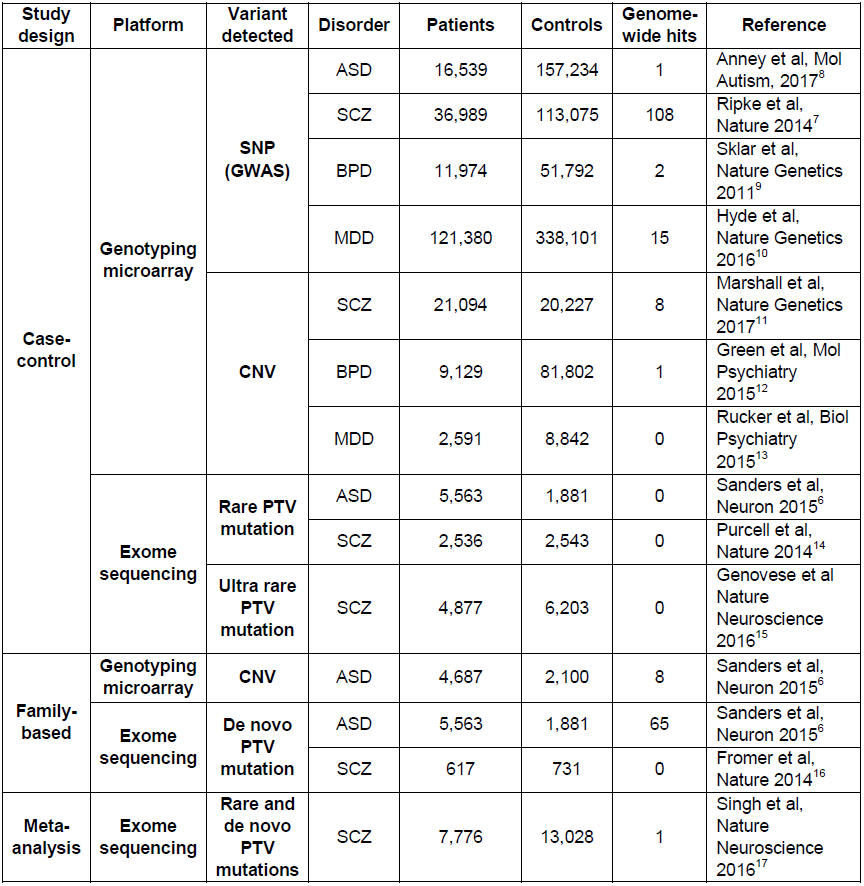
The largest genomic studies to date in autism spectrum disorder, schizophrenia, bipolar disorder, and major depression. SNP, single nucleotide polymorphism; CNV, copy number variant; PTV, protein-truncating variant. ASD, autism spectrum disorder; SCZ, schizophrenia; BPD, bipolar disorder; MDD, major depressive disorder

Exome sequencing, which identifies genetic variants in the ∼1% of the genome that encodes proteins, has identified over 50 genes in ASD (Table 1). The majority of this discovery was through *de novo* protein truncating variants (PTVs) observed in a patient but not in either unaffected parent. Such mutations are very rare, e.g. population frequency ≤0.000002%, but they can have large effect sizes, up to a ∼50-fold increase in risk. As with common variation, these very rare variants have advanced our understanding of the etiology of these disorders, for example by implicating chromatin remodeling in ASD^4,5^.

Although much remains to be discovered, these results have yielded critical starting points for studies of pathogenesis,^6,7^ and indicate the feasibility and importance of discovering sufficient additional variation to fully delineate the key biological pathways underlying these disorders.

### Insights from whole genome sequencing

By assaying most of the genome at single nucleotide resolution, WGS holds the potential to extend rare variant discovery to the ∼99% of the genome that is noncoding (Box 1). While GWAS identifies common noncoding variants, the rare noncoding variants assayed by WGS might have substantially higher effect sizes^1^, increasing tractability for biological experimentation. WGS also enables detection of most structural variation including translocations, inversions, and copy number variants (CNVs)^18,19^. Furthermore, WGS can improve detection of common variants in existing GWAS by statistically inferring SNPs not directly genotyped (imputation) and identifying the specific risk variants within a risk region (fine mapping). Similarly, WGS data may allow detection of common structural variants, including CNVs, that can be missed by current SNP-based approaches^20^, facilitating common CNV association studies.

#### Box 1: Types of genetic variation reliably detected by genomic technologies

**Karyotype (≤ 1% common; ≤ 1% rare):** Chromosomal aneuploidies, massive structural variation (e.g. translocations, inversions, CNVs of millions of nucleotides), some fragile sites with special protocols.

**Microarray (∼90% common; ∼1% rare):** Protein coding and noncoding common SNVs, large rare CNVs (over ∼20,000 nucleotides).

**Exome sequencing (∼1% common; ∼1% rare):** Protein coding common SNVs and indels, protein coding rare SNVs and indels, some CNVs.

**Low coverage WGS (∼95% common; ∼85% rare):** Protein coding and noncoding common SNVs, most protein coding and noncoding rare SNVs.

**Deep coverage WGS (∼99% common; ∼99% rare):** Protein coding and noncoding common SNVs and indels, protein coding and noncoding rare SNVs and indels, rare and common CNVs (over ∼1,000 nucleotides), multi-allelic CNVs (e.g. over 3 copies), mobile element insertions, other structural variation (e.g. translocations, inversions)

**Long-read (>10,000bp), deep coverage WGS (100% common; 100% rare):** As for deep coverage WGS plus: small CNVs (50-1,000 nucleotides), complex structural variation, variants in repetitive DNA, direct assessment of phasing (whether two variants are on the same allele)

SNV: Single nucleotide variant

Indel: Insertion/Deletion (gain or loss of ≤50bp)

CNV: Copy number variant (gain or loss of >50bp).

### The role of noncoding variation

There is considerable evidence that noncoding variation influences brain function and neuropsychiatric disorders. Over 90% of disease-associated GWAS loci discovered by assaying common variants map to noncoding regions^21,22^. In humans, at least 4% of the noncoding genome has been under strong purifying selection^23^. Additionally, epigenomic studies have identified many functional noncoding elements involved in regulation of gene expression underlying neurogenesis, cell differentiation, and neurodevelopment^24^.

Noncoding variation influences which exons are expressed within a gene, in which cells, and under what circumstances. While such insights can be gained from gene association^25^, noncoding variation studies should increase the resolution of such analyses by identifying regulatory regions of genes restricted to fewer cell types, developmental periods, or brain regions. Given the multiple biological roles (pleiotropy) of genes implicated in psychiatric disorders, such WGS-derived hypotheses may be critical for biological follow-up.

### The role of rare noncoding variation

While common noncoding variation clearly plays a role in neuropsychiatric disorders, the role of rare noncoding variation is less clear. A pessimist could note that in Mendelian disorders few linkage peaks were resolved to noncoding causal variants and that systematic deletion of noncoding regions proximate to the *HPRT1* gene (Lesch–Nyhan syndrome) had little impact on protein activity^26^. In contrast, an optimist could argue that Fragile X, the first psychiatric linkage peak resolved to a gene, is a triplet repeat expansion in the 5` untranslated region (UTR) of the *FMRP* gene, and that there are several clear examples of Mendelian traits (e.g. *OCA2* enhancer in eye color) and disorders (e.g. *TBX5* enhancer in congenital heart disease) with penetrant noncoding variants^27^.

The role and utility of rare variation in the noncoding genome is likely to be a function of the number of noncoding regions that, when mutated, disrupt gene expression or function to a high degree. While this can be estimated in model systems, there will be experimental confounds (e.g. species, cell type, developmental stage) that limit interpretation. Direct analysis of WGS offers a complementary and irreplaceable approach to identify and characterize the role of rare noncoding variants in human disease.

WGS technology is sufficiently novel that we cannot accurately evaluate its potential in neuropsychiatric disorders without generating pilot data in human cohorts. It may implicate novel biological pathways missed by previous genomic efforts and identify disease-associated regulatory elements specific to certain cell types, developmental stages, or brain regions. Alternatively, WGS may prove less efficient than cheaper methods in identifying experimentally actionable disease-associated variation. Optimal allocation of future resources rests on efforts, such as the WGSPD, that seriously test the utility of WGS.

### Estimating our ability to find rare noncoding variants

Finding disease-associated loci or variants by WGS will prove more challenging than with GWAS or WES. With WGS there are two orders of magnitude more sites to consider (∼3 billion) compared to potential loci in GWAS (∼20 million) or variants in WES (∼30 million). Furthermore, we cannot predict functional changes, e.g., to transcriptional rate, in the straightforward way we can predict changes to amino acids from coding variation.

To evaluate our power to detect noncoding variants in WGS data, we estimated the power to detect *de novo* protein truncating variants that contribute to risk in ASD^4,28,29^ if they were in the noncoding genome. Without any additional information to help us distinguish signal from noise, for every one risk-mediating variant in the WGS data there would be about 25,000 non-risk variants (a ratio of 1:25000, Table S2). By only considering variants with some evidence of functional effect (e.g. conservation) or proximity to a gene with genome-wide significant association to ASD, we would expect to reduce the noise of non-risk variants, making the risk-mediating variant signal easier to detect. We considered a range of annotation scenarios, from an optimistic 1:5 to a pessimistic 1:500 (see Table S2). Moreover, we do not know what penetrance to expect for these noncoding variants so we considered a wide range, shown as relative risk. For context, the highest relative risks for common variants and *de novo* mutations in psychiatric disorders are about 1.3 and 50, respectively.

We first considered our ability to detect an overall excess of noncoding variants between cases and controls (a burden analysis). Such an analysis could identify a class of variants that mediate risk in psychiatric disorders, for example promoters in proximity to ASD-associated genes, providing insight into regions of the noncoding genome most likely to yield specific risk variants for neuropsychiatric disorders. Since there is no clear category of noncoding variation equivalent to *de novo* protein truncating variants, we adjusted for testing 1,000 annotation categories. The results for *de novo* and case-control analyses are shown in Figure 1a and 1b respectively (see Supplemental Methods).

**Figure 1.**
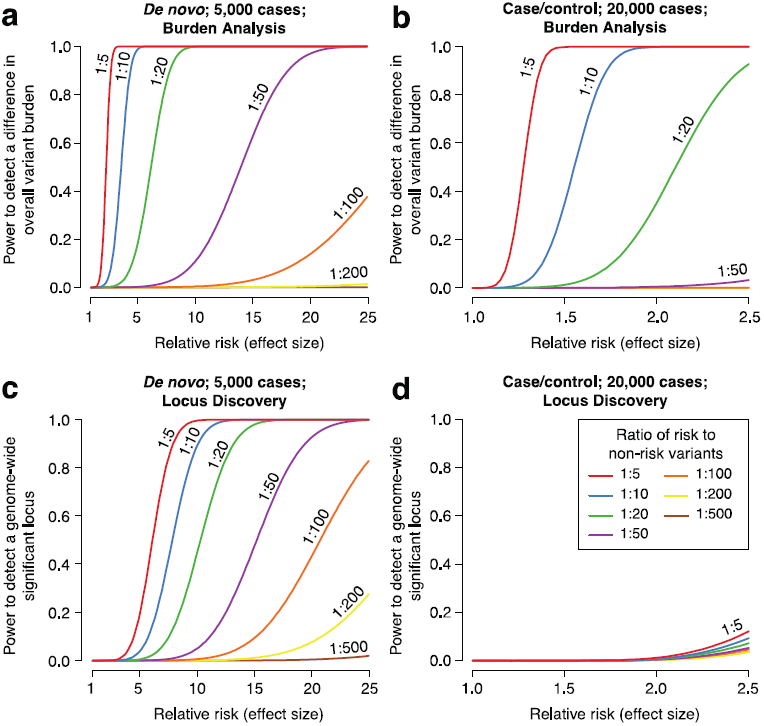
Statistical power in the noncoding genome. We estimated the power at a significance threshold (alpha) of 5 × 10^−5^, to account for 1,000 categories of noncoding variants, to detect an excess of 122,500 noncoding variants in cases vs. controls as we varied the relative risk and risk:non-risk ratio, which represents annotation quality (Table S2). In **a)** we assessed the power for detecting an excess of de novo mutations in 5,000 cases vs. 5,000 controls as the relative risk increases. With a risk:non-risk ratio of 1:20, approximately equivalent to assessing protein truncating variants in the coding genome, we achieve >80% power with a relative risk of 5. In **b)** the power to detect an excess burden of rare variants (allele frequency ≤0.1%) is assessed in 20,000 cases vs. 20,000 controls. In **c)** we assessed the power to identify an excess of de novo mutations at a specific genomic locus, e.g. the noncoding region regulating a single gene. Consequently, we set the significance threshold (alpha) at 2.5×10^−6^. In **d)** we assessed the power to identify an excess of rare variants (allele frequency ≤0.1%) at a specific nucleotide (alpha = 3.3×10^-11^), since this yielded better power than testing for burden at a locus (alpha = 2.5×10^-6^).

We next considered our ability to identify a specific genetic variant, functional element, or group of functional elements (e.g. enhancers that regulate one gene) associated with risk that could be assessed in larger patient cohorts. The results for *de novo* and case-control analyses are shown in Figure 1c and 1d respectively (see Supplemental Methods).

From these analyses, it is clear that we will need: 1) large cohorts, and 2) methods to decrease background noise (to obtain a high risk to non-risk ratio), e.g. through predicting functional effects or regulation of known risk loci.

### Why perform WGS in psychiatric disorders?

Given current uncertainty over the utility of WGS, we could wait until WGS for non-psychiatric phenotypes provide sufficient insight to enable better power analyses. However, even large case-control cohorts may not be informative of the utility of WGS in ASD, for which *de novo* mutations have provided a more efficient approach to identifying specific genes and genetic loci^6,30^ (Figure 1). Additionally, there is a pressing need to identify specific cell types, tissues, and developmental stages involved in brain-based disorders due to the complexity of the nervous system, limited understanding of how molecular changes lead to disorder, and difficulty in interpreting model systems. In short, the potential benefits of WGS in psychiatric disorders may be greater than in other phenotypes and the availability of family-based cohorts may offer insights otherwise unobtainable.

### Implications for neuroscientists

Interpreting the biology downstream of variants identified by existing WES and GWAS analyses remains a challenge; this is especially true in neuroscience due to the inaccessibility and complexity of neural tissue.

The interface of human genetics and neuroscience has typically focused on rare, highly penetrant variants that permit generation of transgenic animals with a robust phenotype^5,31–34^. Neuroscientists now face the challenge of obtaining biological insights through investigation of the multiple weakly penetrant variants, identified through modern genomics, that act through unknown neurological mechanisms, in a manner highly dependent on genetic background^35^.Noncoding variants will pose yet harder challenges. Their effect sizes are likely to be small, and the relevant biology likely to be restricted to specific cell types, developmental stages, or cell states. Analysis of 3D chromatin structure must often be performed to identify the genes that a noncoding variant regulates. Finally, a proportion of noncoding variants may have human-specific functions absent in model organisms. For example, human accelerated regions (HARs), which are conserved across multiple species but differ within humans, are enriched for homozygous variants in consanguineous ASD cases^36^.

Notwithstanding such challenges, many variants identified by genomic technologies have strong evidence of association with the disorders, creating a foundation for investigating pathogenesis. Furthermore, the presence of numerous variants allows systems analyses that identify biological convergences^5^, thus generating mechanistic hypotheses.

### Strategies to improve locus discovery in WGS

#### Sample selection

As with other genomic technologies, **large sample sizes** will be key (see Figure 1 and S2); the simplest way to achieve large cohorts will be through case-control studies, see Table 2.

**Table 1.** Individuals with WGS data generated by, or accessible to, the WGSPD. ^*^ASD samples are being generated by several groups: Centers for Common Disease Genomics (CCDG) of the National Human Genome Research Institute (NHGRI), Simons Foundation Autism Research Initiative (SFARI)^73^, Autism Sequencing Consortium (ASC)^74^.†6,100 samples are shared between Trans-Omics for Precision Medicine (TOPMed) of the National Heart, Lung, and Blood Institute (NHLBI) and CCDG, therefore the total number of samples was reduced by 3,050 for each cohort. These cohorts are composed of individuals ascertained for non-psychiatric disorders and for whom their psychiatric disorder status is generally unknown.

| Data being generated by the WGSPD |  |  |  |  |
| --- | --- | --- | --- | --- |
| Project | Disorder | Cases | Controls | Details |
| 1 | Schizophrenia | 3,333 | 1,667 | Case-control analysis; African American ancestry |
| 1 | Bipolar Disorder | 3,333 | 1,667 | Case-control analysis; African American ancestry |
| 2 | ASD | 378 | 1,512 | Simplex families with two parents, affected child, unaffected child |
| 2 | Schizophrenia | 281 | 843 | Families with two parents and one or more affected children |
| 3 | Schizophrenia | 1,000 | 1,400 | Case-control analysis of individuals from Finland |
| 3 | Bipolar Disorder | 1,000 | 500 | Case-control analysis of individuals from Finland |
| 3 | Schizophrenia | 650 | 325 | Case-control analysis of individuals from Netherlands |
| 3 | Bipolar Disorder | 650 | 325 | Case-control analysis of individuals from Netherlands |
| 3 | Bipolar Disorder | 62 | 138 | Multiplex families with affected and unaffected from Colombia |
| 3 | Bipolar Disorder | 83 | 170 | Multiplex families with affected and unaffected from Costa Rica |
| 4 | Schizophrenia | 271 | 280 | Multiplex families with affected and unaffected |
| 4 | Bipolar Disorder | 299 | 309 | Multiplex families with affected and unaffected |
| 4 | Major depression | 476 | 492 | Multiplex families with affected and unaffected |
| Data being generated by other funding mechanisms with consistent analysis pipelines |  |  |  |  |
|  | Disorder | Cases | Controls | Details |
|  | ASD* | 5,302 | 15,856 | Families with two parents, affected child, +/- unaffected child |
|  | ASD* | 150 | 150 | Multiplex families with affected and unaffected |
|  | Schizophrenia | 118 | 198 | Multiplex families with affected and unaffected |
|  | Bipolar Disorder | 118 | 198 | Multiplex families with affected and unaffected |
|  | Major depression | 478 | 804 | Multiplex families with affected and unaffected |
|  | TOPMed<sup>†</sup> | 0 | 68,950 | Heart, lung, blood and sleep disorders |
|  | CCDG<sup>†</sup> | 0 | 63,950 | Heart, vascular, lung, bowel, neurological, and endocrine disorders |
|  | **Totals** | **17,957** | **165,834** |  |

Several recent studies have shown an excess of deleterious variants in **isolated populations** that have expanded rapidly following recent bottlenecks^37–40^, including deletions of the *TOP3B* gene, associated with schizophrenia and intellectual disability^39^, in ∼3% of individuals in Northern Finland compared to 0.05% in other European populations. **Large multiplex pedigrees** with multiple affected individuals may be enriched for rare, inherited variants with high effect sizes^41,42^. **Simplex pedigrees**, with only one affected individual, are enriched for *de novo* mutations with very high effect sizes given the lack of exposure to natural selection. This strategy has succeeded in severe early-onset disorders, including ID and ASD^4,6,29,43^. Finally, **consanguineous pedigrees** may be enriched for homozygous variants that, like *de novo* mutations, are extremely rare with very high relative risks^36,44,45^. Homozygous variants may also play a role in non-consanguineous cases (Table S4) and have been found to contribute to risk in some outbred ASD families^46,47^. Determining which of these sample selection strategies will be most successful will require WGS pilot projects under each strategy.

#### Integrating phenotypic data

Broadly, two contrasting approaches have been employed in integrating phenotypes in genomic studies, both with the aim of improving statistical power: 1) Combining clinically- or genetically-related diagnoses to increase sample size; and 2) Subdividing cohorts by shared phenotypes to decrease heterogeneity of the underlying genetics (subtyping). GWAS data demonstrate substantial common variant sharing across current conventional diagnostic categories, e.g., bipolar disorder and schizophrenia^48^. Similarly, genes identified by *de novo* mutations are frequently shared between ASD, intellectual disability, and developmental delay^4,29^. Thus, combining data from related diagnoses, can increase sample size, hastening variant discovery^9^.

The alternative approach, subtyping phenotypes, was critical for discovery of Mendelian disorders by linkage methods, in which mis-classifying one individual could prevent discovery. However, such an approach is risky for common, non-Mendelian psychiatric disorders given: 1) current lack of insight into relevant subtypes; and 2) reduced sample size. A GWAS based on ∼2,500 cases in the Simons Simplex Collection ASD cohort showed no improvement in the proportion of genetic heritability explained by the top SNPs accounting for changes in sample size for over 10 phenotypic characteristics^49^. In contrast, a GWAS of a nonpsychiatric phenotype, bone mineral density, showed benefits of subgrouping, leading to the identification of 16 new loci^50^.

Phenotypic subtyping also poses practical challenges. Genetic analysis is comparatively cheap, while deep phenotyping is cumbersome and costly, effectively diminishing sample size. The relative ease of using pre-existing cohorts and registries to inexpensively boost sample size has favored “phenotype-light” sample collection. This balance could be shifted by the adoption of consistent phenotyping schema^51,52^, identification of reliable neuropsychiatric biomarkers, or utilization of electronic medical records. Several large-scale initiatives are already working in this direction, for example deCODE^53^, UK biobank^54^, Geisinger^55^, and the All of Us Research Program (formerly the Precision Medicine Initiative).

#### Identifying functional variants

Our assessment of statistical power (Figure 1) shows that distinguishing variants that are likely to be functional and risk-mediating (i.e. high risk to non-risk ratio) will maximize discovery of specific noncoding variants. Several strategies might help.

##### Annotating the noncoding genome

Annotations may predict functional variants, including: 1) **Conservation** of DNA sequence across species; 2) Regions of **open chromatin**, where DNA is exposed allowing proteins to bind (detected by DNase-Seq or ATAC-Seq); 3) Regions of **active chromatin**, where epigenetic marks suggest transcription of a nearby gene (detected by ChIP-Seq); 4) **Transcription factor binding sites** (detected by ChIP-Seq); and 5) Predicting the **regulatory gene target** using proximity to the variant (<40% accurate^56^) or physical interactions with target loci (e.g. ChIA-Pet) or genome-wide (e.g. Hi-C, 5C)^56^. Of note, many of these annotations may be tissue and developmental stage specific^57–60^.

Large-scale endeavors such as ENCODE^61^ and the Roadmap Epigenome Consortium (REC)^62^ have created a reference for human epigenome annotation. Parallel efforts focused on brain tissue, such as the PsychENCODE Consortium^63^, will help extend these resources^64^.

##### Cataloguing human variation

Building a database of human variation has proven invaluable in interpreting the coding genome^65^ and the Genome Aggregation Database (gnomAD, http://gnomad.broadinstitute.org) extends this approach to WGS. Such data can be used to estimate regions of constraint, (with less variation than expected), suggesting functionality^66–68^.

##### Regions associated with psychiatric disorders

GWAS and WES have defined specific regions of the genome that contribute to psychiatric disorders, particularly in ASD^4,6,29^ and schizophrenia^7^. It is plausible that noncoding variation in proximity to these regions will be enriched for risk-mediating variants.

##### Large variants

On average, large variants, especially deletions, have greater potential to mediate risk than small variants^6^. However, while large indels and small CNVs may have a greater impact on noncoding function, there are considerably fewer such variants compared to SNVs^18^. The utility of this strategy will depend on the balance between these two opposing effects.

##### Functional validation

Methods have been developed to assess the functional effects of large numbers of potential regulatory regions. These Massively Parallel Reporter Assays (MPRA)^69^, including Self-Transcribing Active Regulatory Region Sequencing (STARR-Seq)^70^, assess the function of a regulatory region by its potential to transcribe itself, or a specific sequence of DNA (barcode). Of note, this ability to functionally validate noncoding variants *en masse* is a major benefit over interpreting coding missense variants, for which protein-specific functional assays are usually required.

### The Whole Genome Sequencing Consortium for Psychiatric Disorders (WGSPD)

The potential for WGS to help understand neuropsychiatric disorders, and the absence of insight into the role of rare noncoding variants, prompted the United States National Institute of Mental Health (NIMH) to fund four pilot projects aimed at generating WGS data in neuropsychiatric disorders to provide a more complete understanding of genomic architecture.

Big questions in biology are akin to solving problems of similar complexity in other disciplines such as particle physics or astronomy and require a ‘Team Science’ approach^71^. Recognizing the need for large samples sizes to make progress (Table 1, Figure 1), the NIMH, the Stanley Center for Psychiatric Research, and researchers at 11 academic institutions across the USA that were funded in the four selected projects, have formed a public-private partnership: the Whole Genome Sequencing Consortium for Psychiatric Disorders (WGSPD). This consortium aims to establish a repository of WGS data, processed in a consistent manner, to facilitate large-scale analyses within and across four psychiatric disorders (Figure 2). This approach can make more efficient use of funding and resources, for example, by using a central data repository, consistent analysis pipelines, and collaborative methods development to help all researchers access and use the data.

**Figure 2.**
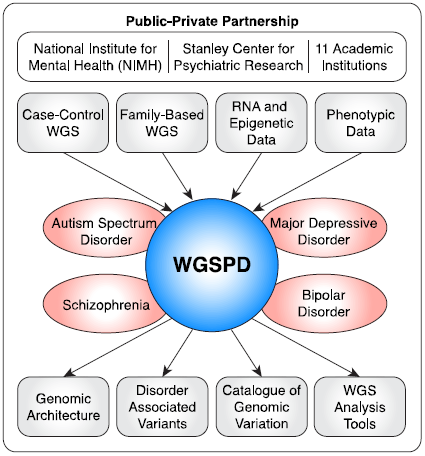
Overview of the WGSPD.

The WGSPD will need to expand, both beyond the founding members and these four disorders. Investigators with relevant WGS data will be invited to join the WGSPD and participate in working groups focused on specific disorders or cross-disorder projects. Given the scale of WGS data, the cost of reprocessing the data in a consistent manner and storing the data will be substantial. Establishing a suitable funding strategy for such genomic integration is a key question that needs to be addressed urgently throughout the genomics community. In a first step to improve this, WGS analysis pipelines have been coordinated across several major sequencing centers and consortia (e.g. CCDG, TOPMed, WGSPD) to allow direct comparison of results. To obtain the sample sizes necessary (Figure 1), a similar consensus will need to be established internationally.

### Cloud-based analysis

The sheer scale of WGS datasets necessitates new models for data analysis, since data storage and computation is likely to be beyond the resources at any single institution. Fortunately, the development of cloud-based computing has coincided with the generation of WGS data. Under this model, a single cloud-based data repository can be accessed by teams at each collaborating site, and cloud-based analysis eliminates the need for cumbersome and costly downloads. This approach has the further advantage of facilitating the sharing of preinstalled algorithms and pipelines, encouraging consistent consortium-wide analysis.

The scale of WGS data can make simple analytical tasks overwhelming. Therefore, the WGSPD is committed to developing Application Program Interfaces (APIs) and software solutions for the wider community to simplify cloud-based data access (e.g. hail^72^). In doing so, computational biologists and analysts can focus on the development and application of methods for analysis, rather than on lower level data management and handling.

The analysis of deidentified genetic data on university-hosted remote servers is common practice, with contributing sites being responsible for securing non-genetic identifying information. So long as cloud environments meet equivalent security standards to existing remote servers, then existing informed consent will cover this use, except in rare instances where the consent specifically excludes this approach. Best practice guidelines for secure sharing of genomic data have been described by the NIH: https://www.ncbi.nlm.nih.gov/projects/gap/pdf/dbgap_2b_security_procedures.pdf. There is an urgent need for methods that allow such guidelines to be easily adopted and readily vetted across cloud providers and institutions.

### The WGSPD projects and data

The four WGSPD projects, developed by independent sets of investigators, encompass the diverse strategies for improving locus discovery and therefore will provide some of the earliest opportunities to assess their relative utility in complex disorders. The four projects are:

1. Case-control analysis of schizophrenia and bipolar disorder in individuals of African American ancestry.
2. Family-based analysis of ASD in families with a single affected child, allowing the detection of *de novo* mutations.
3. Case-control analysis of schizophrenia or bipolar disorder in isolated populations with recent population bottlenecks.
4. Family-based analysis of schizophrenia, bipolar disorder, or major depression in families with multiple affected individuals.

Combining these WGS cohorts with consistently processed WGS data from other consortia will yield an initial dataset of 183,000 individuals, including 18,000 cases and 165,000 controls (Table 1). In addition to the genotype data, we are collating phenotype data that are comparable across projects, disorders, and ages to allow in-depth genotype-phenotype analysis.

## Conclusion

The noncoding genome remains largely unexplored and major discoveries undoubtedly await intrepid explorers. Whole genome sequencing of neuropsychiatric cases and controls provides an important avenue in this exploration, potentially offering high resolution insight into the developmental stages, brain regions, cell types, and biological functions that underlie these disorders. If the cost of sequencing continues to fall, it is inevitable that WGS will ultimately replace both microarray and WES – the key question is at what price point this transition offers a good return for investment. Pooling preliminary WGS data between researchers and across disorders offers the most efficient mechanism to make this determination.

The creation of the WGSPD has allowed numerous researchers to pursue diverse scientific approaches on multiple psychiatric disorders, while simultaneously working towards a harmonized data set for integrated analysis. The pooling of expertise, methods, and data will accelerate progress towards understanding genetic contributions to brain development, function, and pathology and create a resource that will continue to yield scientific and clinical insights for years to come.

## Acknowledgements

The authors would like to acknowledge and thank the study participants and their families. The WGSPD is a public-private partnership between the National Institute of Mental Health (NIMH), the Stanley Center for Psychiatric Research, and researchers at 11 academic institutions across the USA. This work was supported by grants from the National Institute of Mental Health (NIMH): U01 MH105653 (M.B.), U01 MH105641 (S.A.M.), U01 MH105573 (C.N.P.), U01 MH105670 (D.B.G.), U01 MH105575 (M.W.S., A.J.W.), U01 MH105669 (M.J.D., K.E.), U01 MH105575 (N.B.F., D.H.G., R.A.O.), U01 MH105666 (A.P.), U01 MH105630 (D.C.G.), U01 MH105632 (J.B.), U01 MH105634 (R.E.G), U01 MH100239-03S1 (M.W.S., S.J.S., A.J.W.), R01 MH095454 (N.B.F.), the Simons Foundation: (SFARI #385110, M.W.S., S.J.S., A.J.W., D.B.G., SFARI #401457 (D.H.G)), and a gift from the Stanley Foundation (S.E.H.).

## Author contributions

S.J.S., B.M.N., H.H., and D.M.W. contributed the power calculation. All authors contributed to the text.

